# A large single-participant fMRI dataset for probing brain responses to naturalistic stimuli in space and time

**DOI:** 10.1101/687681

**Authors:** K. Seeliger, R. P. Sommers, U. Güçlü, S. E. Bosch, M. A. J. van Gerven

**Author notes:** Corresponding author: K. Seeliger. Equal contribution (K. Seeliger, R. P. Sommers).

## Abstract

Visual and auditory representations in the human brain have been studied with encoding, decoding and reconstruction models. Representations from convolutional neural networks have been used as explanatory models for these stimulus-induced hierarchical brain activations. However, none of the fMRI datasets currently available has adequate amounts of data for sufficiently sampling their representations. We recorded a densely sampled large fMRI dataset (TR=700 ms) in a single individual exposed to spatiotemporal visual and auditory naturalistic stimuli (30 episodes of BBC’s *Doctor Who*). The data consists of 120.830 whole-brain volumes (approx. 23 h) of single-presentation data (full episodes, training set) and 1.178 volumes (11 min) of repeated narrative short episodes (test set, 22 repetitions), recorded with fixation over a period of six months. This rich dataset can be used widely to study the way the brain represents audiovisual input across its sensory hierarchies.

## Background & Summary

Group neuroimaging studies have proven useful for studying effects that have strong similarity across individuals, and that generalize across the population. However, group analysis requires adjustments to the data that have to be made to enable analysis across diverse individuals, which includes the smoothing of functional data and the transforming of brains of different individuals so that they fit a template brain. Additionally, due to the often small amount of data and the differences in data quality stemming from variation in psychophysiological conditions, group studies are not ideal for many questions in sensory neuroscience. Especially questions associated with hypothesized maps of sensory representations require the sampling of large feature spaces. Therefore, large single-participant fMRI datasets seem a good addition to group studies, as it can be more useful to have a dense recording of the various stages in sensory processing in a single healthy brain rather than shorter and lower-SNR recordings of many participants [22, 21, 24].

In addition, fMRI analyses in the encoding model framework are typically done at voxel resolution by building individual models for each voxel to map their function [27]. The regularized linear regression models typically used for encoding models need relatively large datasets. Several single-participant datasets suitable for encoding are currently available to the community. Researchers have started acquiring high-quality general-purpose sensory perception fMRI datasets with the goal of making them publicly available for encoding and decoding analysis [12, 6]. However, for analysing complex sensory representations (for instance the hierarchies in convolutional neural networks) encoding models with massive amounts of free parameters are necessary. Currently available datasets are not sufficient in size for training them adequately.

With the data collected in the present study we aim to take a leap forward for single-participant functional data. We provide a (by current neuroscientific standards) massive dataset of functional data recorded from one individual. We recorded 120.830 volumes of training data and 1.178 volumes of test data (22 repetitions) over the course of six months. Choosing BBC’s *Doctor Who* [8], we provide data with engaging spatio-temporal visual and auditory stimulation with high visual and semantic variety. To ensure positional stability across sessions our set-up made use of several measures, including a head cast [26]. The acquisition was done with fixation on the centre of the screen.

The targeted domain of this dataset is the field of research concerned with the neural coding of naturalistic stimuli via encoding models, representational similarity, multivariate decoding and reconstruction. Scene and emotion processing, semantic representations and scene boundaries in relation to hippocampal activity are further vectors of scientific investigation. Moreover, the prolonged experiment, lasting six months, allows investigating whether there are fine-grained changes in functional responses or functional connectivity over this time.

Cognitive neuroscience has benefited from advances in pattern analysis for a few decades. By providing a dataset of this size, we aim to push the boundaries for applying state-of-the-art machine learning and neural networks methods for understanding sensory processing.

## Methods

### Experiment design

Data recording was done on a single participant (male, age 27.5) between April 2017 and December 2017, with preceding pilot sessions starting in December 2016. In every session, one full *Doctor Who* episode was presented. In most sessions, this episode was followed by a presentation of the seven test set clips in random order. At the end of many of the sessions, a structural scan was acquired, which we optionally omitted depending on the fatigue of the participant and on the scanner schedule.

The design of each session is illustrated in Figure 1. Episodes were split up into four parts, where the first three were exactly 12 minutes long and the last part had variable length, depending on the duration of the respective episode. Stimulus videos were buffered before beginning the MRI recording and started when the fifth MRI volume pulse was received. Each coherent fMRI training set recording presented two parts concatenated to each other, with a very brief time interval in between to wait for the next scanner pulse. Every first video part (i.e. in one fMRI recording consisting of two parts) started with a consistent delay of 100 ms due to buffering, which may be taken into account during analysis. Each part of two was followed by a black screen with the centred word FIN, lasting 16 seconds, during which we continued to collect fMRI volumes to account for the haemodynamic delay (fadeout screen).

**Figure 1:**
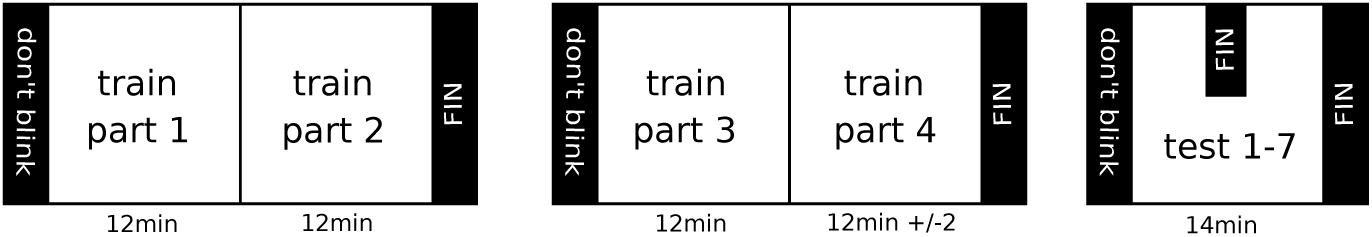
Session design. Every episode was split into four parts with three parts of length 12 minutes and one of variable length. Two parts were presented in each coherent fMRI recording at regular playback speed, followed by a fade-out black screen lasting 16 seconds. The test set was recorded following most sessions, with test set videos in random order and each followed by the fade-out screen.

We ensured that participant positioning was stable, that motion parameters within and between sessions were kept to a minimum and that the participant was comfortable during the experimental procedure. The final set-up was decided upon within ten piloting sessions. Episodes used during this piloting phase were not reused. These piloting sessions were additionally used by the participant for practising staying in the scanner with little motion for a long time, and fixating while viewing videos with strongly varying visual centres of attention.

During the piloting phase we observed a slow gradual head displacement in z-direction during the first 15 minutes of each functional scan, most likely related to changing properties of the magnetic field due to warming (known as frequency drift). After this initial phase, the z-direction displacement plateaued. We accounted for this by a 13-minute functional MRI scan directly after the participant had been placed in the scanner, in which we made an additional presentation of the test set (data not used but available upon request). During this phase we also tested audio quality and adjusted volume together with the participant. After this phase, we proceeded with gradient field map estimation and the functional scans. Breaks between recording blocks were kept short due to the frequency drift, which did not impair the participant’s self-reported attention. Data for these additional test set recordings is available upon request. The fMRI scanning protocol is listed in Table 1.

**Table 1:** fMRI scanning protocol for regular recording sessions.

| Item | Duration |
| --- | --- |
| MRI localizer | 60 s |
| Warm up | 13 min |
| AAScout | 60 s |
| GRE field map | 120 s |
| Training set parts 1 and 2 | 24 min |
| Training set parts 3 and 4 | 24 min +/- 2 min |
| Test set | 13 min |
| T1 anatomical scan | 5 min |

### Ethical statement

Data collection was approved by the local ethical review board (NL45659.091.14, CMO region Arnhem-Nijmegen, The Netherlands, CMO code 2014-288) and was carried out in accordance with the approved guidelines. An amendment of the general ethical approval was acquired due to the extraordinary length of this experiment. Informed consent was given by the participant for every separate session.

### Session fMRI protocol

The participant made sure to remain naive towards the content of every episode prior to the session. We provide a table called *session notes* that associates the file containing the functional data with each episode ID. It also documents the order of the test set videos every session. Likewise, any divergence from the standard protocol as well as special physiological conditions such as tiredness are documented in this table.

### Creating the stimuli

We presented the videos at 20 horizontal and 20 vertical degrees of the visual field. To this end, we undertook a number of resizing and cropping steps on the videos. We resized the videos to 698 × 1264 in order to reduce the amount of the picture that would be discarded. Then they were cropped to near-square (732 × 698) without further reducing resolution. The near-square resolution was needed for correcting the beamer projection via the mirror in our scanner at target visual field coverage. For analysis, the stimuli should be resized to square as a final step, since the participant perceived the stimuli as a square. Stimuli were presented on a mirror reflecting a projection from a beamer outside the MRI system. As a last step, a black margin was added in the cropped regions to reach the beamer display resolution of 1024 × 768. These steps are illustrated in Figure 2.

**Figure 2:**
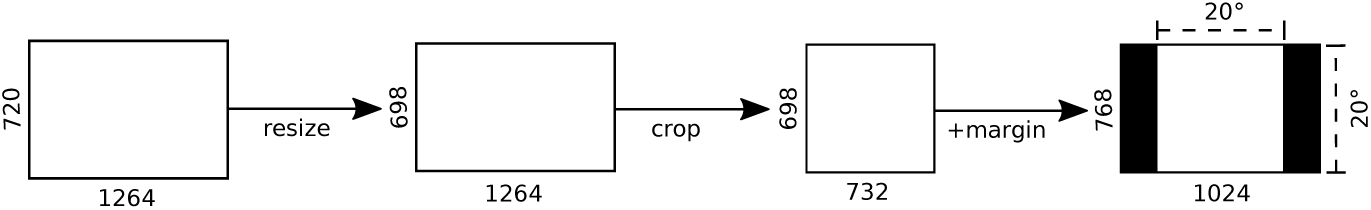
Stimulus preparation. *In order to present the video on 20 degrees of the visual field (horizontally and vertically) we resized and cropped them to squares of size* 732 × 698, *adding black margins to fill up the beamer resolution.*

#### Dynamic range compression

We compressed the dynamic range of the audio signal as the volume of speech and other sounds (e.g. explosions) was overrepresented, leading to speech being overshadowed by the scanner noise at tolerable volumes for the other sounds. `Ffmpeg` and `sox` were used for dynamic range compression as follows (example code for Season 4, Episode 1, Part 1):

~~~
$ ffmpeg -i avi_S04_E01_P01.avi -vn -acodec copy mp3_S04_E01_P01.mp3 sox mp3_S04_E01_P01.mp3 compand_mp3_S04_E01_P01.mp3 compand 0.3,1 6:-70, -69,-30 -20 -90 0.2
$ ffmpeg -i avi_S04_E01_P01.avi -i mp3_S04_E01_P01.mp3 -c:v copy -c:a copy -strict experimental -map 0:v:0 -map 1:a:0 cavi_S04_E01_P01.avi
~~~

#### Video and audio codecs

After splitting up into parts and dynamic range compression, all stimulus video parts were re-encoded from H.264 to the *Indeo 5.1 video codec* and the *PCM raw audio format (uncompressed)*, which are recommended and locally validated codecs for time-accurate use with *Presentation* and the local fMRI stimulus presentation computer. This was done with a `VirtualDub` script (involving the `ffdshow` filters)) which is available from the data repository.

#### Fixation

The participant was instructed to fixate on the centre of the video scene. Train and test videos were thus superimposed with a central cross (coloured cyan, which is least likely to occur in the video stimuli) optimal for fixation [25]. The fixation cross is available in the data repository. Superimposing it may not be necessary for most data analyses. The technical validation presented in this manuscript was done with fixation crosses imposed on the video data.

#### Training video sources and parameters

Training set episodes were extracted from the original BluRays (Season 2: `BBCBD0262`, Season 3: `BBCBD0263`, Season 4: `BBCBD0264`).

Table 2 lists the parameters of the original videos (before splitting, dynamic range compression and then re-encoding) and the actually used stimulus videos. The repository contains a PDF listing additional technical information on the stimulus re-encoding and creation.

**Table 2:** Training video parameters of original video files and of stimulus videos.

| Parameter | Original videos | Stimulus videos |
| --- | --- | --- |
| Container format | .mkv | .avi |
| Video codec | H.264 | Indeo 5.1 |
| Resolution | 1264x720 | 732x698 |
| Frame rate | 23.976215 Hz | 23.976023 Hz |
| Audio codec | DTS Audio | PCM uncompressed |
| Audio sampling rate | 48 kHz | 44.1 kHz |

**Table 3:** Information about video clips used in the test set.

|  | Episode | Title | Test ID | Repetitions | Length |
| --- | --- | --- | --- | --- | --- |
| Pond Life | 1 | April | 1 | 26 | 60 s |
|  | 2 | May | 2 | 26 | 60 s |
|  | 3 | June | 3 | 26 | 60 s |
|  | 4 | July | 4 | 26 | 60 s |
|  | 5 | August | 5 | 26 | 60 s |
| Space / Time | 1 | Space | 6 | 22 | 180 s |
|  | 2 | Time | 7 | 22 | 180 s |

#### Stimulus presentation

Stimuli were projected on a rear-projection screen using an Eiki projector with a resolution of 1024 × 768, placed outside the shielded room. A stimulus PC with the Presentation software package was used to display the stimuli.

#### Training stimuli

For the training set we used episodes from Seasons 2, 3 and 4 after the 2005 relaunch of the series (*Tenth Doctor*). We started with Episode 6 from Season 2 and presented all following episodes in serial order until and including Season 4, Episode 10. As mentioned previously, every episode was split up into four parts before any dynamic range compression or re-encoding. This was done using `ffmpeg`as follows:

~~~
$ ffmpeg -t 12:00.000 -i mkv_S02_E06.mkv -vcodec h264 avi_S02_E06_P01
$ ffmpeg -ss 12:00.001 -i mkv_S02_E06.mkv -vcodec h264 -t 12:00.000 avi_S02_E06_P02.avi
$ ffmpeg -ss 24:00.001 -i mkv_S02_E06.mkv -vcodec h264 -t 12:00.000 avi_S02_E06_P03.avi
$ ffmpeg -ss 36:00.001 -i mkv_S02_E06.mkv -vcodec h264 -t 13:37.000 avi_S02_E06_P04.avi
~~~

The end of each episode was determined as the moment when the *Doctor Who* logo appeared. This also determined the length of each fourth part.

#### Test stimuli

The test set consisted of seven variable length videos. We used *Pond Life*, a mini-series of 5 narrative 1-minute-episodes; and *Space / Time*, two mini-episodes of 3 minutes each. They were originally written for later seasons in the series (*Eleventh Doctor*) and feature similar characters, but not the same actors. We decided for these test set stimuli since they covered high visual and semantic variability in a short time, and because taking later material from later seasons meant that there was no scene overlap. Space / Time was extracted from a DVD `(BBCDVD3430)`. The one-minute Pond Life clips can be obtained from official promotional material.

The test set videos were cropped and re-encoded in the same way as the training set stimuli (see Figure 2 & Table 4) and therefore ended up with the exact same resolution as the training set videos. However, as they were taken from a different source than the training stimuli, they still have slightly different video parameters. One difference between test set videos and training videos is the change in aspect ratio caused by resizing. For test set videos 1-5 the original resolution was 1280 × 720 and its original aspect ratio 1.78. Resizing this to 1264 × 698 and an aspect ratio of 1.81 does not lead to a vastly different *change* in aspect ratio than the training set (from 1.76 to 1.81). Test set videos 6 and 7 were only available in a resolution (720 × 576) with a much smaller aspect ratio (1.25) than the desired one. Resizing them to 1264 × 698 and aspect ratio 1.81 resulted in a greater change in aspect ratio due to resizing than was the case for the other videos. This greater change meant that a slight, yet noticeable elongation of the video in the vertical axis was visible during stimulus presentation. However the change was slight enough that this went unnoticed during the experiment.

**Table 4:** Test video parameters of original video files and of stimulus videos.

| Parameter | Original videos | Stimulus videos |
| --- | --- | --- |
| **Episodes 1,3-5:** |  |  |
| Container format | .mp4 | .avi |
| Video codec | H.264 | Indeo 5.1 |
| Resolution | 1280x720 | 732x698 |
| Frame rate | 25 Hz | 25 Hz |
| Audio codec | MPEG AAC Audio | PCM uncompressed |
| Audio sampling rate | 44.1 kHz | 44.1 kHz |
| **Episode 2:** |  |  |
| Container format | .mp4 | .avi |
| Video codec | H.264 | Indeo 5.1 |
| Resolution | 1280x720 | 732x698 |
| Frame rate | 30.013455 Hz | 30.013455 Hz |
| Audio codec | MPEG AAC Audio | PCM uncompressed |
| Audio sampling rate | 44.1 kHz | 44.1 kHz |
| **Episodes 6-7:** |  |  |
| Container format | .mkv | .avi |
| Video codec | MPEG-1/2 Video | Indeo 5.1 |
| Resolution | 720x576 | 732x698 |
| Frame rate | 25 Hz | 25 Hz |
| Audio codec | A52 Audio | PCM uncompressed |
| Audio sampling rate | 48 kHz | 44.1 kHz |

A second difference between videos is that while the training set has a frame rate of 24 Hz, the frame rate of the test set is 25 Hz - with the exception of test 2, which has a frame rate of 30 Hz. A third difference is that different video and audio codecs were used to encode the video information in the original files. While we converted all videos to the exact same codecs, the conversion process might theoretically create small differences between stimuli.

Initially we regarded these differences between the seven test set videos and the training set as undesirable confounds in our stimuli and debated leaving test videos 2, 6 and 7 out of the data set. However, we decided that the deviations could actually yield additional information about our models. If models that are trained on our training set remain able to predict the test set fMRI responses - despite the above differences in low level features - then that must mean that the model must have learned features invariant to these differences. Therefore, we have decided to leave these slightly different test stimuli in the greater data set and leave it up to the user of our data to decide whether to include or exclude them in their analysis. We do recommend keeping the differences in mind when using our data.

The test set was presented in random order after the training episode. After every video there was a 16 second blank screen allowing us to collect the remaining signal of the hemodynamic response. Table 3 and Table 4 summarize all of the information about the test set and associates each clip with the test set ID used in the data records.

### fMRI data acquisition

The data was collected in a Siemens 3T MAGNETOM Prisma with a Siemens 32-channel head coil (Siemens, Erlangen, Germany). The functional scans used a T2*-weighted echo planar imaging pulse sequence at 700 ms TR. Volumes were recorded with 64 transversal slices at 2.4 mm^3^ voxel size (slice dimension 88 × 88 at field of view 211 × 211 mm^2^). They were measured with a multiband acceleration factor of 8. TE was 39 ms and the flip angle 75 degrees. The multiband-multiecho protocol allowed recording from multiple transversal locations simultaneously at high speed, which is beneficial for video stimuli. The structural scans at the end of most sessions used a T1-weighted MP RAGE pulse sequence, with 192 sagittal slices with a field of view of 256 × 256 and 1 mm^3^ voxel size. The TE was 3.03 ms, the TR 2300 ms and the flip angle 8 degrees.

### Positioning

To ensure stable positioning across sessions, a custom-made 3D-printed foam head cast [26] was fitted for the participant and the MRI coil. Positioning in the z-direction was further stabilized with a polymer chin rest. Additionally, before each session we checked participant position by measuring the distance between the participant’s nasion and the edge of the head cast, and the distance between the head cast and both pupils. As the head coil mount of the MRI scanner model has (stable) leeway, we also ensured that it was in the same position for each experiment.

Comfortable positions of neck cushions, elbow cushions, shoulder bags and a knee cushion were determined during the piloting phase until motion parameters were minimal, and then kept constant throughout the experiment. The soft thick mattresses of the MRI table were replaced with a thin mattress on top of a wooden plank to increase position stability and reduce neck strain.

As a last step during positioning, we ensured that video presentation was at a constant location in the visual field. We did this by using a physical horizontal line attached to the projection mirror that the participant had to match to a projected horizontal line by adjusting the mirror tilt inside the scanner before the session. The lines were removed before starting the experiment.

Note that the procedure we describe and none of the existing ones can account for movement or deformation of the brain inside the skull.

#### Further consistent preparations

Audio was presented via S14 Insert Earphones by Sensimetrics. The volume was adjusted to functional scanning conditions (with scanner noise, during warm up) prior to the sessions to ensure that the participant could effortlessly follow the narration. Since caffeine has a measurable effect on the BOLD response [7], the participant kept his caffeine intake constant. He also had to cut his hair in regular intervals as thickening hair gradually decreased comfort inside the head cast.

### Localizer sessions

In addition to the regular functional data recording sessions, we carried out retinotopy and recorded data for auditory and functional localizers within three separate sessions. Localizers for identified areas with similar specializations partly overlap.

#### Retinotopy

Retinotopy was done with polar wedges, using `FSL v5.0` for analysis. We mapped V1, V2, and V3; separately for left and right hemisphere and dorsal and ventral sections.

#### Functional localizers

We collected a series of functional localizers related to the visual and auditory systems. They were estimated as contrasts with `FSL`. The localizer set and partly the recording procedure from [14] served as inspiration and reference for our localizer choices and acquisition methods. Images were selected from image stimulus databases for faces [17], animals and objects [4, 5], and human bodies without faces [9]. The functional localizers are provided separately for left and right hemisphere, and individually for separate categories if we used multiple for mapping them. The estimated functional area locations were verified by comparing to other studies using `Neurosynth` [28] (using a transformation to MNI space).

LOC, OFA and FFA were identified using recordings from the same session, with 16 blocks of 16 seconds length in which 20 images were shown. Images were displayed for 300 ms with a pause of 500 ms in-between.

#### LOC

The *lateral occipital complex* was identified with contrasts of animals, faceless bodies, everyday objects against scrambled versions of the same categories.

#### FFA

The *fusiform face area* [15, 19] was identified with contrasts of human faces, animals and faceless bodies against scrambled versions of the same categories.

#### OFA

The *occipital face area* was identified with contrasts of human faces against scrambled faces.

We localized AC, M1 and MT within separate sessions.

#### AC

The *auditory cortex* was identified with contrasts of audio stimuli versus resting state activity. We used a block design consisting of 10 blocks with three different conditions. Blocks were presented for 20 seconds (the duration of the stimuli), and consisted of a fixation cross and audio file. We contrasted music (the introductory music of the series), the Tardis sound, and speech (the main character of the series talking) with rest in order to map the AC.

#### M1

For identifying the *primary motor cortex* we asked the participant to make small movements for the duration of the cue presentation. This was done using six distinct block types randomly distributed over 30 blocks. Each block was presented for 16 seconds, with a 12 second inter block interval. In the Rest condition, we only presented the word REST, the participant was told not to move during this block. For the Hand and Foot blocks, we presented the words HAND and FOOT, after which the participant had to move his left and right *hand* (finger drumming movements) and *feet* (toe movements) respectively. For the Mouth block, the word MOUTH was presented and the participant produced nonsensical syllables involving lips and tongue for (*mouth*) movement. For the *speech* localizer, SPEECH was presented and he was asked to sub-vocalize sensible sentences. Finally, for the Eyes block, we only presented 9 dots in a 3 × 3 grid. The participant was asked to make random saccades in quick succession to each dot throughout the grid. We contrasted the motion blocks with rest in order to localize the M1.

#### MT

The *middle temporal visual area* [3], sometimes also called V5 was identified using a block design with static dots, coherently moving dots, and rest. In a 5 minute session, we presented 17 blocks, each block lasting for 15 seconds. The blocks with static dots and coherently moving dots consisted of 600 dots that either remained in the same location, or moved with constant speed through the visual display. In the rest block we only presented a fixation cross. We contrasted the moving dot blocks with the static and rest blocks in order to localize MT.

### Preprocessing

We converted the `DICOM` images to `*.nii` using `dcm2niix`^1^. Preprocessing was done using `FSL v5.0` as follows: We realigned the series of volumes recorded for each 12 minutes part (called one *run*) to their middle volume (reference volume). We then estimated the transformation from each run to the middle (reference) volume from the very first run in the series. We applied this transformation to the train and test data, which resulted in volumes realigned to the middle volume of the first run. The preprocessing script is available from the data repository. Note that our preprocessing only consists of realignment and no further steps. Due to our multiband protocol it also does not involve slice time correction.

## Data Records

The acquired participant data records described in this section are available at the *Donders Repository*^2^, persistent ID `11633/di.dcc.DSC_2018.00082_134`^3^. Additional information such as information about how to recreate the stimuli and their exact length is available from there as well. As we only applied light preprocessing (realignment) and as we aim for the data to be ready-to-use (with comparable results) we provide the train and test fMRI recordings in preprocessed format, and with volumes recorded before the start of the stimulus video discarded.

The acquired participant data records described in this section are available at the (*anonymous data repository)*, persistent ID (anonymous DOI)^4^. Additional information such as information about how to recreate the stimuli and their exact length is available from there as well. As we only applied light preprocessing (realignment) and as we aim for the data to be ready-to-use (with comparable results) we provide the train and test fMRI recordings in preprocessed format, and with volumes recorded before the start of the stimulus video discarded.

### Train fMRI responses

Available are 120 fMRI recordings in 4D NIfTI format of exactly (part 1-3 of every episode) or approximately (part 4) 12 minutes length. At a TR of 700 ms and a 14 seconds fadeout screen, this has lead to 1049 volumes per 12 minutes recording. In total there are 120.830 training set volumes including fadeout and 119.630 volumes when discarding the fadeout volumes. The file name format for the training data is `rrun[001-121].nii.gz` and they are located in `func/train`. Note that there is no data for `rrun[005]` since it was omitted from public release due to a known video buffering problem during presentation. For internal bookkeeping the original numbering of the data was kept intact. `rrun[005]` was not omitted for the analyses presented in the technical validation.

### Test fMRI responses

Recordings of 22 repetitions of 7 narrative test set videos, provided in 4D NIfTI format and located in `func/testavg` (averaged and standardized with zero mean and unit variance) and `func/test` (individual presentations). For the recordings of the more visually varied test set videos 1 to 5, four additional recordings are available (26 in total). The individual presentations use the file structure `repetition[repetition number]_run_[test video ID].nii.gz`. The averaged presentations are contained in a `Matlab` file (readable by `python scipy.io` and `Matlab`) with key name scheme `mean[test video ID]`. In total 25.927 volumes were recorded including fadeout, and 27.032 when not counting fadeout volumes. The data is available in single presentation and in averaged format. The test set averaged over all repetitions is the one intended to be used for analysis. The length of this averaged section is 1.178 including the fadeout volumes. Test sets were recorded in most, but not all sessions. Refer to the session notes for further information.

### Structural data

We provide the anatomical T1 reference scan in the directory `anat`. The scan has been defaced with `mri_deface` for anonymisation.

### Localizers

Retinotopic and functional localizers are available as 88 × 88 × 64 masks in NIfTI format. Functional localizers were acquired as contrasts.

### Session notes

The root directory contains `train_session_notes.csv` and `test_session_notes.csv` that include information about every session. There is information on which episode was shown, special physiological circumstances, whether a test set was recorded (and in which test video order), and which episode was shown.

### Information for stimulus creation

The directory `stim/preparation` contains detailed information and scripts for recreating the Doctor Who episodes we used as stimuli. It also contains the fixation cross `(fixation_cross.bmp)`.

### Additional scripts

The directory `code`/ contains an `FSL v5.0` script for preprocessing the data. Our procedure is described in the respective methods section.

## Technical Validation

### Motion across sessions

We analysed the stability of our set-up by determining the translation and rotation parameters via motion correction with FSL `mcflirt`. Figure 3 presents the motion parameters relative to the first training set volume recorded in `run001`. This figure shows to what extent we were able to recreate the target initial head position before the start of every session using our positioning method. While the translation in the x direction was stable across sessions, in the z and y directions we observed translations, with a maximum of 2 mm in z direction over all sessions. The mean translation of y and z was 0.6 mm and 0.88 mm respectively (all parameters ignoring one outlier session addressed below, which had 8 mm translation). The mean pitch and roll rotation (yaw was stable) was 0.45° in each direction, with a max rotation of 2° in roll direction. With a voxel size of 2.4 mm^3^ this is certainly not as optimal as it could be, but still within an acceptable range for effective realignment across 30 sessions.

**Figure 3:**
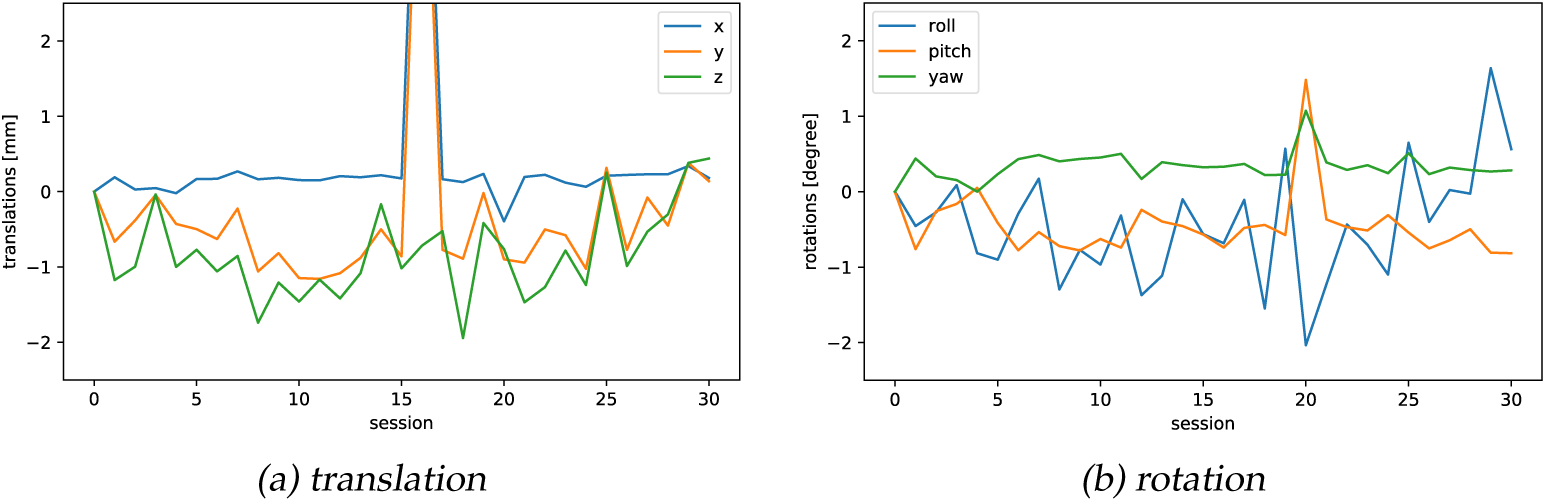
Motion parameters across sessions. Translation (3a) and rotation (3b) estimated for each initial training volume in the series with `mcflirt`, relative to the first training volume recorded in the first session. Represents to what extent we were able to recreate the target position before every session with our positioning method. The session numbers on the x axis are continuous starting from 0 (session SES03) and do not match the session IDs.

The outlier session in the series had an x translation of 7.2 mm (session number 16 in the figures, which had session ID SES20 and run numbers 62-65). The participant reported pressure pain for the second fMRI training run and the test set, which however also occurred in few further runs without such translation problems. We assume that during this session we failed to verify that the head coil base was in the right position (it has 1 cm leeway), but are unable to confirm it at this point. Our preprocessing procedure did not fail realigning this session and (after realignment) the data seemed usable and was not excluded for further analysis. However, excluding this session from analysis should be considered in case it leads to problems.

Figure 4 presents the same motion parameters *within* every training session, with the first training volume recorded in every session as reference. These figures indicate how well the participant could keep his position stable across the session. For about half of the sessions we notice a ramp up, similar to the frequency drift effects described previously. The dip in the trajectory is the gap between the two fMRI recordings runs (each consisting of two video parts). The persisting, yet lower frequency drift-related motion overshadowed the magnitude of the participant’s actual motion parameters. We conclude from this that the participant was able to keep his head position mostly stable within a session.

**Figure 4:**
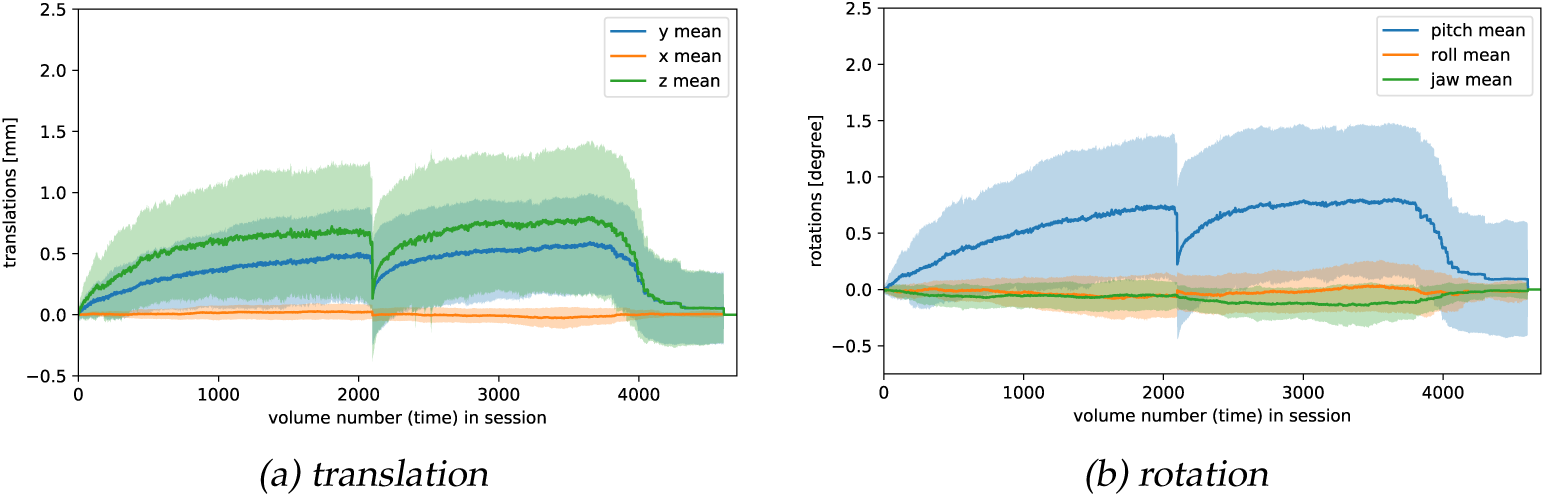
Motion parameters within sessions. Mean translation (4a) and rotation (4b) parameters within the training sessions. They are estimated with `mcflirt`, relative to the first volume recorded in every training session. The error bars show the standard deviation. This represents how well our participant could keep his initial position stable across a session. Since his movements were smaller than the frequency drift effects, the latter overshadow the participant’s movements.

### Motion energy visual encoding model

Motion energy features [1] are a powerful representation of spatiotemporal visual data. Using voxel-wise encoding models it was shown that motion energy features based on a spatiotemporal Gabor filter pyramid can model BOLD activity in wide sections of the occipital lobe responding to natural videos [20]. *Voxel-wise encoding* means that linear regression models predicting BOLD data based on stimulus information are trained and evaluated independently for every voxel.

For quality assessment we evaluated the voxel-wise motion energy encoding model with our data. We limited the analysis to cortical voxels. As the code for motion energy feature extraction for fMRI video stimuli has been made available by the authors^5^ we could stay close to the original feature extraction method. It resulted in 6555 features, which were averaged over the TR of 700 ms. For accommodating the haemodynamic delay we shifted stimulus matrices temporarily in a step-wise fashion and concatenated the steps four times so that every feature could potentially influence the voxel response recorded in a volume from 2.8 s to 5.6 s after the stimulus event. Using multiple possible peak delay times will also account for small differences in the haemodynamic delay across the brain. This procedure is described in detail in the supplementary material of Nishimoto 2011 [20]. For every voxel the full time-shifted and concatenated motion energy matrices present the input to a regression model. The regression models are regularized using the L2 norm (ridge regression). The regression models learn to predict the responses of single voxels at one point in time, given a time-shifted and concatenated feature matrix for that time point. They are trained on the single presentation training data. Predictive performance was quantified using the Pearson correlation between actual and predicted responses. The resulting voxel-wise correlations on our dataset are projected on the brain and can be seen in Figure 5. For every voxel-associated model the optimal regularization parameter *λ* is determined by taking the best *λ* from the average correlation over 10 folds of 1% randomly selected training data points. Models are trained and evaluated on detrended and standardized voxel responses (separately for every fMRI recording) and standardized motion energy features. We excluded the fadeout periods – due to this and the FIR-like analysis around 35 volumes per training run are not used, leading to 23 h of data left to analyze. We projected the voxel-wise correlations onto a flatmap using pycortex [10]^6^ in combination with `FreeSurver v6.0.0` and `FSL`. The result is shown in Figure 5 It indeed confirms that early and intermediate visual areas can be explained well with the motion energy model on our dataset.

**Figure 5:**
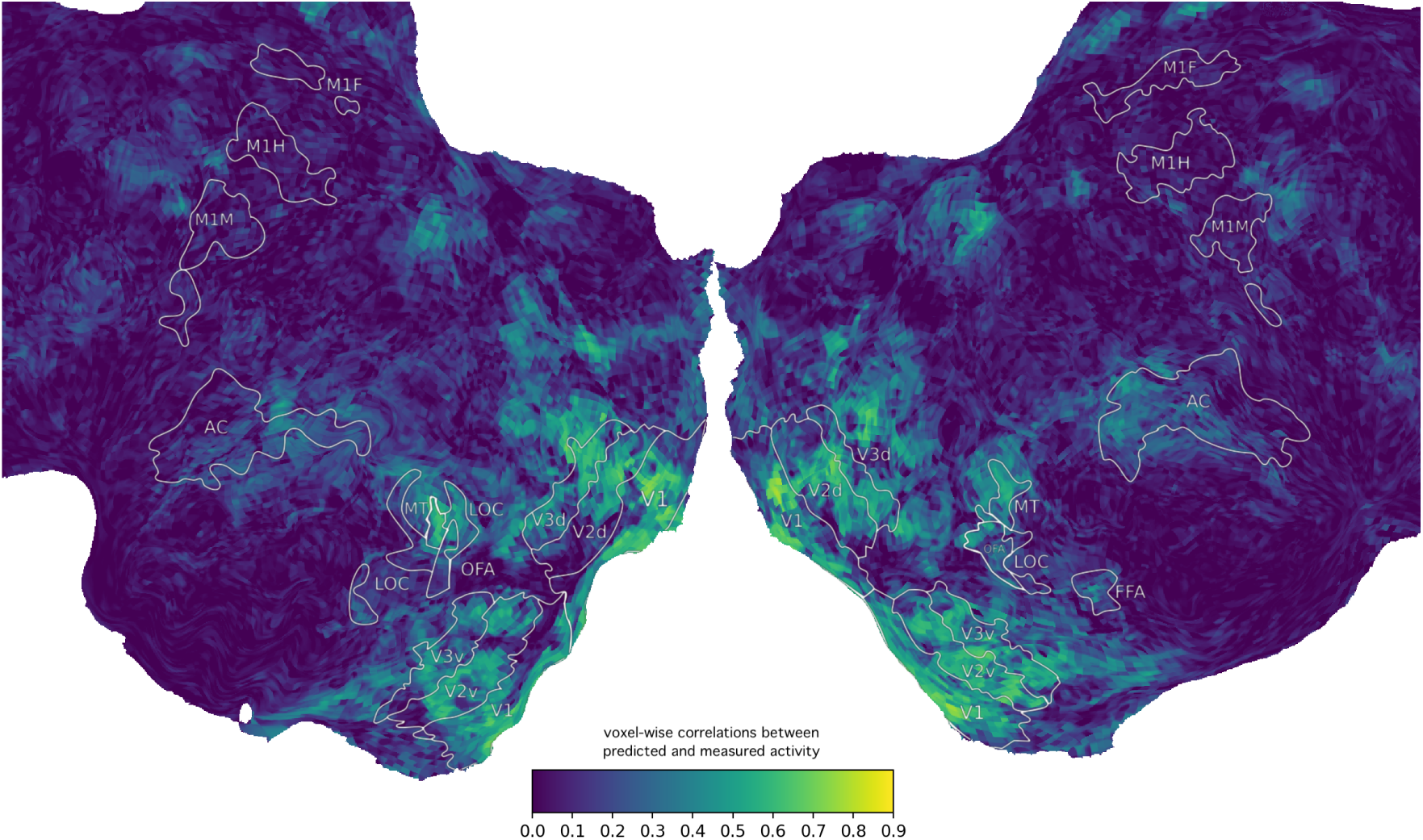
Voxel-wise correlations of the motion energy encoding model. The color map encodes Pearson’s correlations between the measured BOLD activity and the activity predicted by each linear regression model on the held-out test set.

We also looked into how the performance of the motion energy encoding model behaves as a function of the amount of training data used. The mean correlation of significant voxels over the number of training hours can be found in Figure 6, evaluated per session.

**Figure 6:**
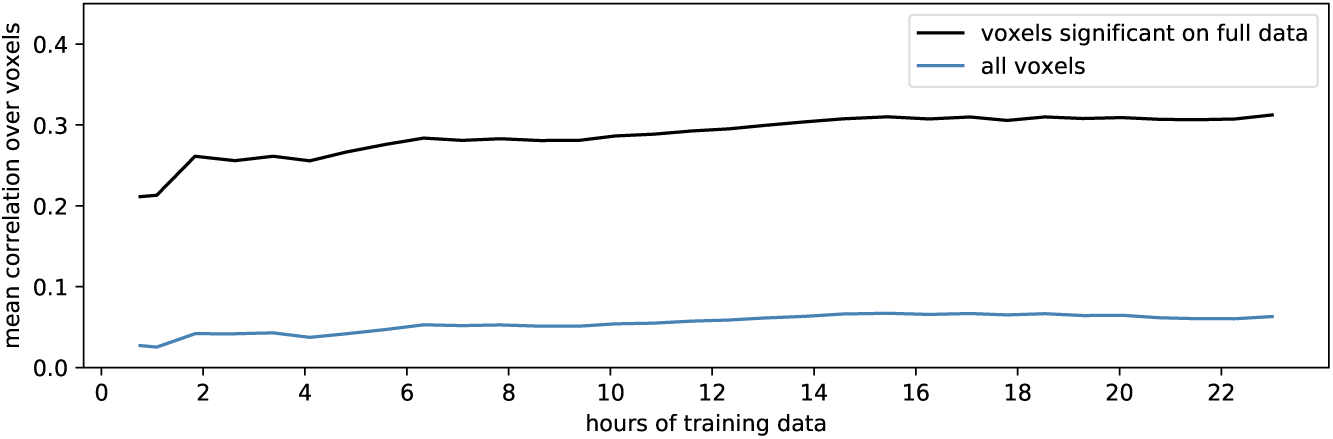
Average correlation across significant voxels. (p<0.01, Bonferroni-corrected over number of voxels) over amount of training data. The selection of significant voxels was done on the training run with maximum data.

This average correlation of motion energy encoding models can also be studied separately for every ROI. We assume that in the early visual system maximum correlations will be reached faster, while in higher visual cortex regions they will need more training data to reach a plateau. We indeed observe this behaviour for our participant in Figure 7, where visual cortex areas V1, V2, V3 and MT reach the maximum performance after two hours of training data while the higher order areas we localized, especially FFA only seems to peak at more than 16 hours of training samples from our participant. This observation is probably characteristic of the motion energy encoding model which is a hypothesized representation for early visual system areas, but may also affect models that cover higher order areas such as convolutional neural networks.

**Figure 7:**
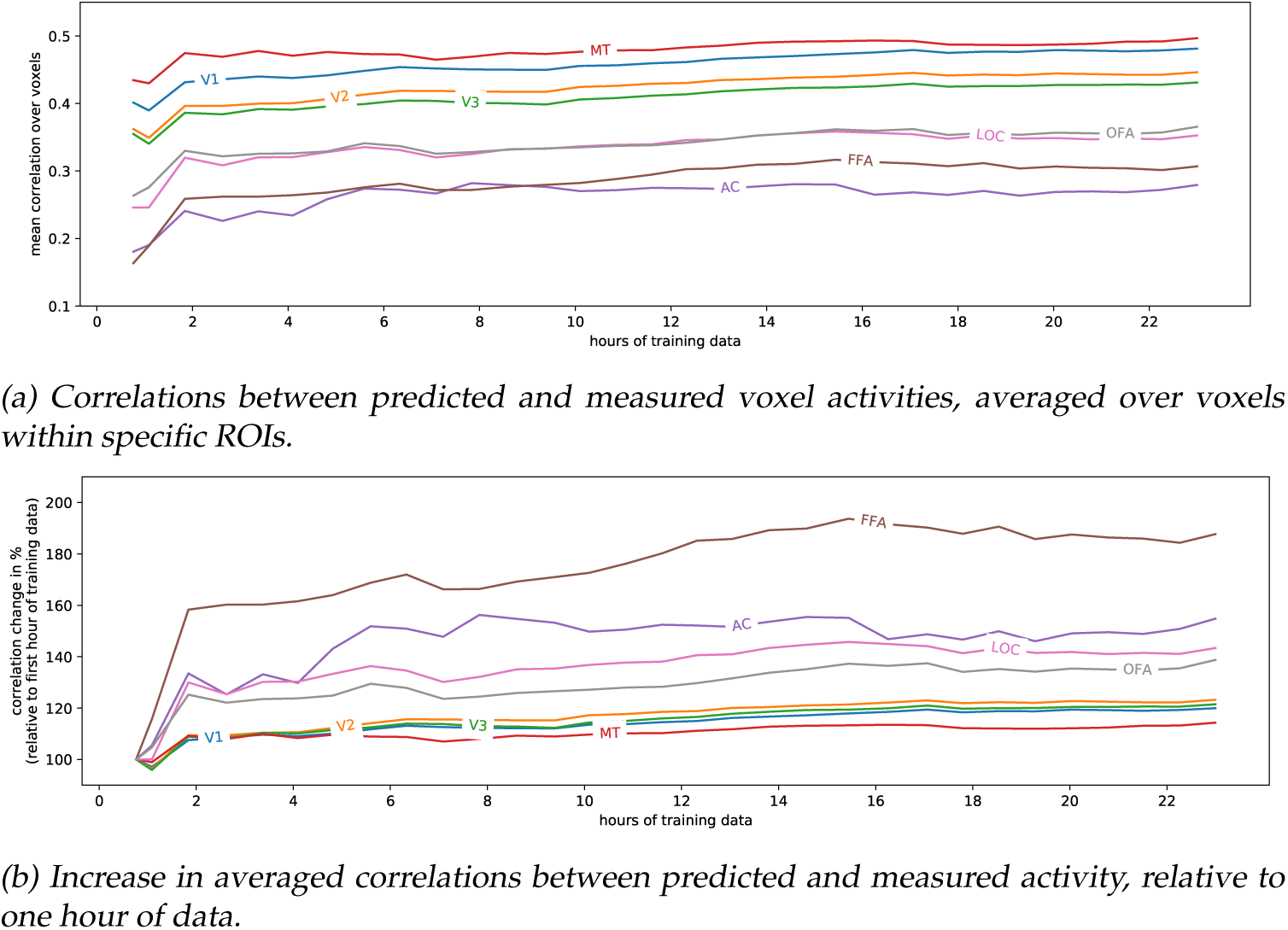
Average correlation across significant voxels in specific ROIs. (p<0.01, Bonferroni-corrected over total number of voxels) over amount of training data. 7a shows average correlations for significant voxels with increasing amounts of training data, starting from the first session. 7b shows ROI-wise average correlations relative to using only one (the first) hour of training data. We see that the early visual cortex reach their peak correlation much faster than higher order areas.

Figure 8 shows average correlations over significant and all voxels estimated for individual sessions. While there are differences between sessions, they appear to stay in the same range and we did not observe severe outlier sessions in terms of performance with the motion energy model.

**Figure 8:**
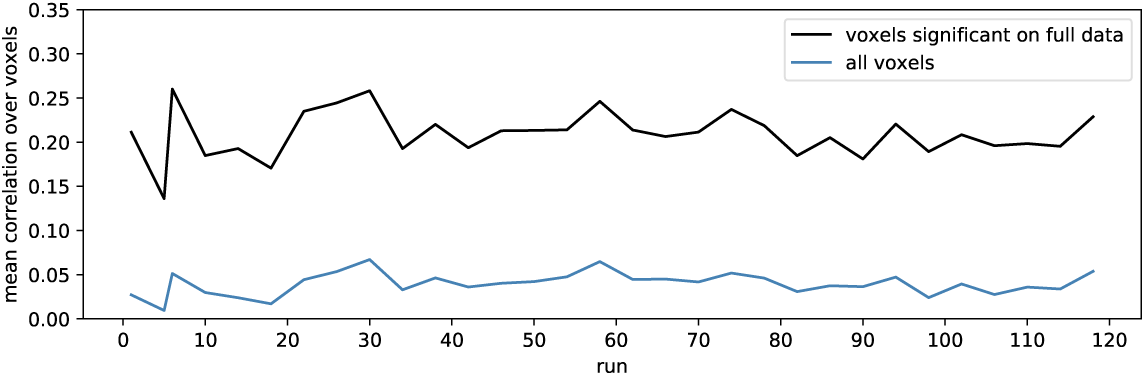
Average correlation across significant voxels. (p<0.01, Bonferroni-corrected over number of voxels) for individual sessions (specified by their run numbers). The selection of significant voxels was again done on all data.

## Usage Notes

Please cite this data descriptor when using the data set for research purposes. See section *Data Citations* at the end of this document for an example citation. The original authors of this data set do not have to be listed as co-authors in subsequent research using it. For legal reasons related to our data use agreement `RU-DI-HD-1.0` researchers who want to access the data have to reveal their identity to the study authors when requesting access via *Donders Repository*.

## Acknowledgements

We would like to express our gratitude to Paul Gaalman for help with developing the fMRI sequence and with stabilizing the experimental set-up. We would also like to thank Sam Lawrence and David Richter for help with localizer development.

This research was supported by VIDI grant number 639.072.513 of The Netherlands Organization for Scientific Research (NWO).

## Author contributions

RS, UG, KS, SB and MvG designed and piloted the experiment. RS created the stimuli, experiment code and localizer experiments and organized the data acquisition. UG and SB designed the fMRI protocol and preprocessed the data. SB analysed the localizers. RS, SB, KS and UG acquired the data. KS analysed the data for technical validation. KS prepared the data for analysis and publication. KS wrote the data description manuscript. KS, SB, MvG and RS finalized the manuscript.

## Data citations

This dataset is published exclusively for research purposes. Please use the following citation when using this data for your own research:

Seeliger, K., Sommers, R. P., Güçlü, U., Bosch, S. E., van Gerven, M. A. J. (2018): *A large single-participant fMRI dataset for probing brain responses to naturalistic stimuli in space and time*. Dataset published on *Donders Data Repository*. Persistent identifier: `11633/di.dcc.DSC_2018.00082_134`.

## Footnotes

1 `dcm2niix from 28th March 2018`

2 `data.donders.ru.nl`

3 Direct link to the repository: http://hdl.handle.net/11633/di.dcc.DSC_2018.00082_134

4 Direct link to the repository: (direct link removed due to anonymization)

5 github.com/gallantlab/motion_energy_matlab (accessed December 2017)

6 `pycortex` from June 30th 2018

